# Evolution of an intratumoral ecology susceptible to successive treatment in breast cancer xenografts

**DOI:** 10.1101/249797

**Authors:** Hyunsoo Kim, Pooja Kumar, Francesca Menghi, Javad Noorbakhsh, Eliza Cerveira, Mallory Ryan, Qihui Zhu, Guruprasad Ananda, Joshy George, Henry C. Chen, Susan Mockus, Chengsheng Zhang, Yan Yang, James Keck, R. Krishna Murthy Karuturi, Carol J. Bult, Charles Lee, Edison T. Liu, Jeffrey H. Chuang

**Author notes:** **Corresponding Author:** Jeffrey Chuang. The Jackson Laboratory for Genomic Medicine, 10 Discovery Drive, Farmington CT 06032.

## Abstract

The processes by which tumors evolve are essential to the efficacy of treatment, but quantitative understanding of intratumoral dynamics has been limited. Although intratumoral heterogeneity is common, quantification of evolution is difficult from clinical samples because treatment replicates cannot be performed and because matched serial samples are infrequently available. To circumvent these problems we derived and assayed large sets of human triple-negative breast cancer xenografts and cell cultures from two patients, including 86 xenografts from cyclophosphamide, doxorubicin, cisplatin, docetaxel, or vehicle treatment cohorts as well as 45 related cell cultures. We assayed these samples via exome-seq and/or high-resolution droplet digital PCR, allowing us to distinguish complex therapy-induced selection and drift processes among endogenous cancer subclones with cellularity uncertainty <3%. For one patient, we discovered two predominant subclones that were granularly intermixed in all 48 co-derived xenograft samples. These two subclones exhibited differential chemotherapy sensitivity -- when xenografts were treated with cisplatin for 3 weeks, the post-treatment volume change was proportional to the post-treatment ratio of subclones on a xenograft-to-xenograft basis. A subsequent cohort in which xenografts were treated with cisplatin, allowed a drug holiday, then treated a second time continued to exhibit this proportionality. In contrast, xenografts from other treatment cohorts, spatially dissected xenograft fragments, and cell cultures evolved unsystematically but with substantial population bottlenecks. These results show that ecologies susceptible to successive retreatment can arise spontaneously in breast cancer in spite of a background of irregular subclonal bottlenecks, and our work provides to our knowledge the first quantification of the population genetics of such a system. Intriguingly, in such an ecology the ratio of common subclones is predictive of the state of treatment susceptibility, suggesting that this ratio can be measured to optimize dynamic treatment protocols in patients.

**AUTHOR SUMMARY:** An overarching challenge of cancer is that patients develop resistance to treatment -- an essentially evolutionary process. However, there is currently very little understanding of how tumor evolution can be exploited to improve treatment. One reason for this is that usually only 1-2 samples can be obtained per patient, so cancer evolutionary processes are still poorly understood. To solve this problem, we created many dozens of copies of the tumors from two breast cancer patients using xenografting and cell culture methods. We then compared the evolution in these tumor copies in response to different treatments, including four of the most common breast cancer chemotherapies. These studies present the most exhaustive comparisons of treatment-induced evolution that have yet been performed for individual cancer patients. Unexpectedly, high-resolution sequencing of these samples revealed a special dynamically treatable ecology in one tumor, in which tumor growth during platinum therapy was determined by the ecological balance of two tumor cell populations. Our work shows that ecologies that can be targeted by dynamic treatment strategies arise spontaneously in breast cancers. Population heterogeneity is common within cancers, and our work suggests how tracking of intratumoral evolution can be used to optimize treatment.

## INTRODUCTION

It has long been theorized that differential response of subclones to therapy is important to the development of tumor drug resistance [1–4]. Genomics approaches have now accelerated the ability to study intratumoral population evolution, e.g. as reviewed in [5]. Still, understanding of the dynamics by which tumor subclonal distributions evolve, and in particular in response to therapy, remains limited. An increasingly common approach has been to sequence fragments of patient tumors from multi-region sampling or primary/recurrence/metastasis comparisons [6]. However, regional samples taken synchronously from a patient tumor will not distinguish the effect of treatment, and determining why sequences differ among regional samples is difficult because their relative growth histories are unknown. Comparison of recurrences, metastases, and primaries is similarly limited by the lack of information during the many months that often separate clinical samples [7–9].

Patient-derived xenografts (PDXs) are a model system in which human tumors can be grown in mice for controllable time intervals under specified treatments, providing a solution to these challenges [10–12]. PDXs recapitulate therapy response of patient tumors while also retaining some intratumoral heterogeneity. This makes them a valuable system for studying the evolution of tumor subclonal populations, both as a pool for resistant cells [13] and as an ecology with potentially important interpopulation interactions [14].

Another challenge to understanding tumor heterogeneity is that common measurement protocols such as exome-seq have biases that can influence estimates of allele frequency [15] and hence subclone cellularity levels. Cellularity inference methods are also sensitive to copy number [16], and computational copy number estimates from sequencing or array data are strongly dependent on model assumptions [17]. Due to these complexities, prior heterogeneity studies have usually focused on whether samples share mutations, rather than interpreting quantitative cellularity values [18–20]. In patient-derived xenografts, state-of-the-art heterogeneity studies have used exome-seq and copy number array measurements with Dirichlet process subclone inference algorithms to estimate intratumoral composition, with a main qualitative finding being that there are strong subclonal changes at first engraftment [21,22]. Other groups have measured subclonal evolution in xenografts derived from cell lines [14,23], though their heterogeneity differs from patient-derived populations. Still, elucidation of the evolutionary response during therapy remains a critical challenge, and accurate quantification will be important for designing new treatment strategies [24].

In this study we comparatively investigate how subclonal populations respond to four distinct treatments using patient-derived xenografts for triple negative breast cancers (TNBCs), an aggressive subtype with a high propensity to progress [25]. Using xenografts and conditionally reprogrammed progenitor cells derived from two patients, we finely quantify how the chemotherapies cisplatin, docetaxel, doxorubicin, and cyclophosphamide impact cancer cell populations. As a control, we also investigate the effects of spatial and temporal parameters. In total, we perform sequencing on 131 samples, providing a deep interrogation of the behaviors within these patients’ cancers. To resolve measurement uncertainties, we make extensive use of droplet digital PCR, a high precision technology to determine matched mutations and copy number changes. These data allow us to perform, to our knowledge, the most exhaustive deconstructions to date of the evolution of phenotypically-relevant tumor populations across treatments and biological replicates. Finally, we analyze these results to identify intratumoral ecological behaviors that may be targetable by dynamic treatment strategies.

## RESULTS

### Triple-negative breast cancer xenografts exhibit intratumoral heterogeneity

We established TNBC xenografts from two female breast cancer patients using fragments (5-10 mm^3^) obtained from the patient tumor at resection through engraftment into Nod scid gamma mice (see Methods). Tumors were grown and further dissected over 3-4 passages to generate multiple xenografts from each tumor. To investigate the effect of chemotherapy on tumors and their subclonal populations, for each of the two patient tumors we generated five cohorts of 8-9 xenografts and treated each with one of cisplatin, docetaxel, doxorubicin, cyclophosphamide, or a vehicle control (Figure 1A). For both models, tumors continued to grow during doxorubicin and cyclophosphamide treatment and responded to docetaxel. Tumor shrinkage was observed in response to cisplatin for one model (TM00099) while for the other model (TM00096) a low level of growth was observed.

**Figure 1.**
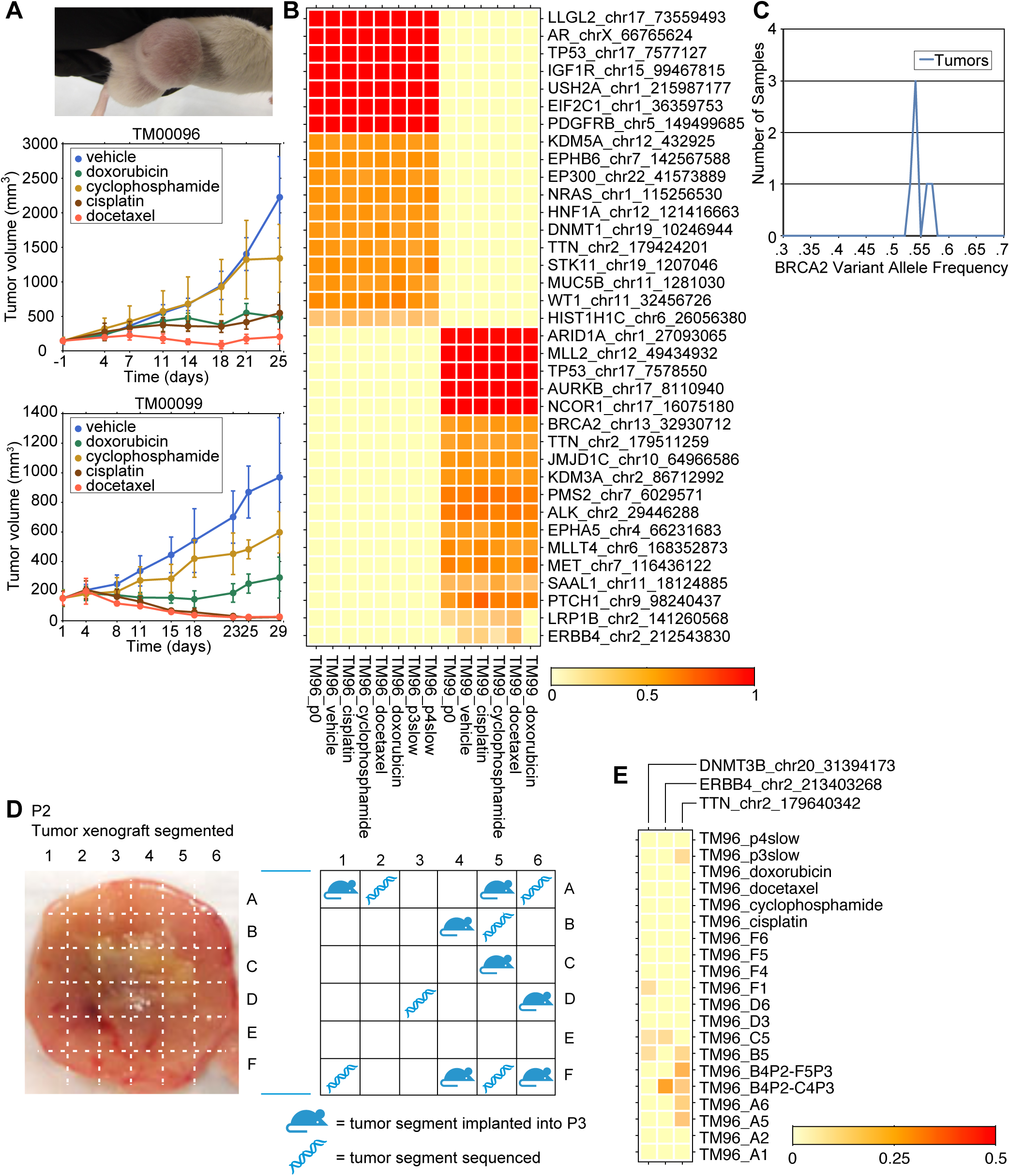
Exome-seq quantifications of mutations in triple-negative breast cancer xenografts. (A) Treatment response curves for cohorts of patient-derived xenografts generated from two triple negative breast cancers (TM00096 and TM00099). For each patient tumor, cohorts of 8 mice each were generated for cisplatin, doxorubicin, cyclophosphamide, docetaxel, and a vehicle control. (B) Heatmap of allele frequencies for point mutations identified in residual tumor samples from treated xenografts. (C) Distribution of BRCA2 mutant allele frequencies for 6 TM00099 xenograft samples as measured by exome-seq cancer panel. The mean and standard deviation of mutant allele frequency are 0.545±0.015. (D) Spatial dissection of the P2 passage of tumor TM00096. (E) Mutations predicted by MuTect to differ among samples in the TM00096 tumor.

To assay subclonal changes, we initially sequenced one residual sample per treatment arm and other untreated samples using an exome-capture platform covering 358 cancer-associated genes at 400× sequencing depth (see Methods). We computationally identified cancer mutations and their allele frequencies (AFs) after filtering out mouse sequence (Figure S1, see Methods). Most recurrent variants exhibited stable AF across all samples and often at values ~50% or >90%, suggesting truncal mutations or germline polymorphisms (Figure 1B). These provided a quantification of exome-seq measurement uncertainty, e.g. a variant in *BRCA2* in the TM00099 tumor yielded AFs of 0.545±0.015 across 6 samples, compared to the AF=0.5 heterozygous expectation (Figure 1C). Some loci exhibited stable AF values at lower levels (e.g. *HIST1H1C* in the TM00096 tumor exhibited 0.18<AF<0.27 across samples) suggesting modulation by copy number (CN) changes, and computational CN estimates qualitatively supported this conclusion (see Methods). Other loci showed sample-specific variation, e.g. an *ERBB4* mutation in the TM00099 tumor was confirmed by Sanger sequencing to be private to certain samples (Figure S2). We also observed AF differences among untreated spatiotemporally-separated samples, which included 6 fragments from a dissected TM00096 xenograft grown for 118 days (Figure 1D, Figure S3) and 7 fragments from further passaged TM00096 xenografts (See Methods), indicating drift processes were non-negligible on the 2 month time scale (Figure 1E). Nevertheless, we found that sample-specific quantifications of subclone cellularities were limited by high uncertainties in CN inferences (Figure S4), consistent with prior studies [26,27].

To resolve these uncertainties in measuring subclones, we used droplet digital PCR sequencing (ddPCR, Figure 2A) to resequence some prior as well as newly generated samples. We chose a mix of loci with stable or variable behavior across samples based on the exome-seq data, performing AF or CN measurements at 23 loci for the TM00099 tumor samples and 18 loci for the TM00096 tumor samples. To better survey conditions, we replicated the treatment experiments on additional xenograft cohorts for the two patient models and sequenced multiple individuals from each cohort while tracking their tumor volumes. We also performed new cell culturing and serial therapy retreatment studies as described below, yielding a total of 1633 ddPCR measurements for either allele frequency or copy number.

**Figure 2.**
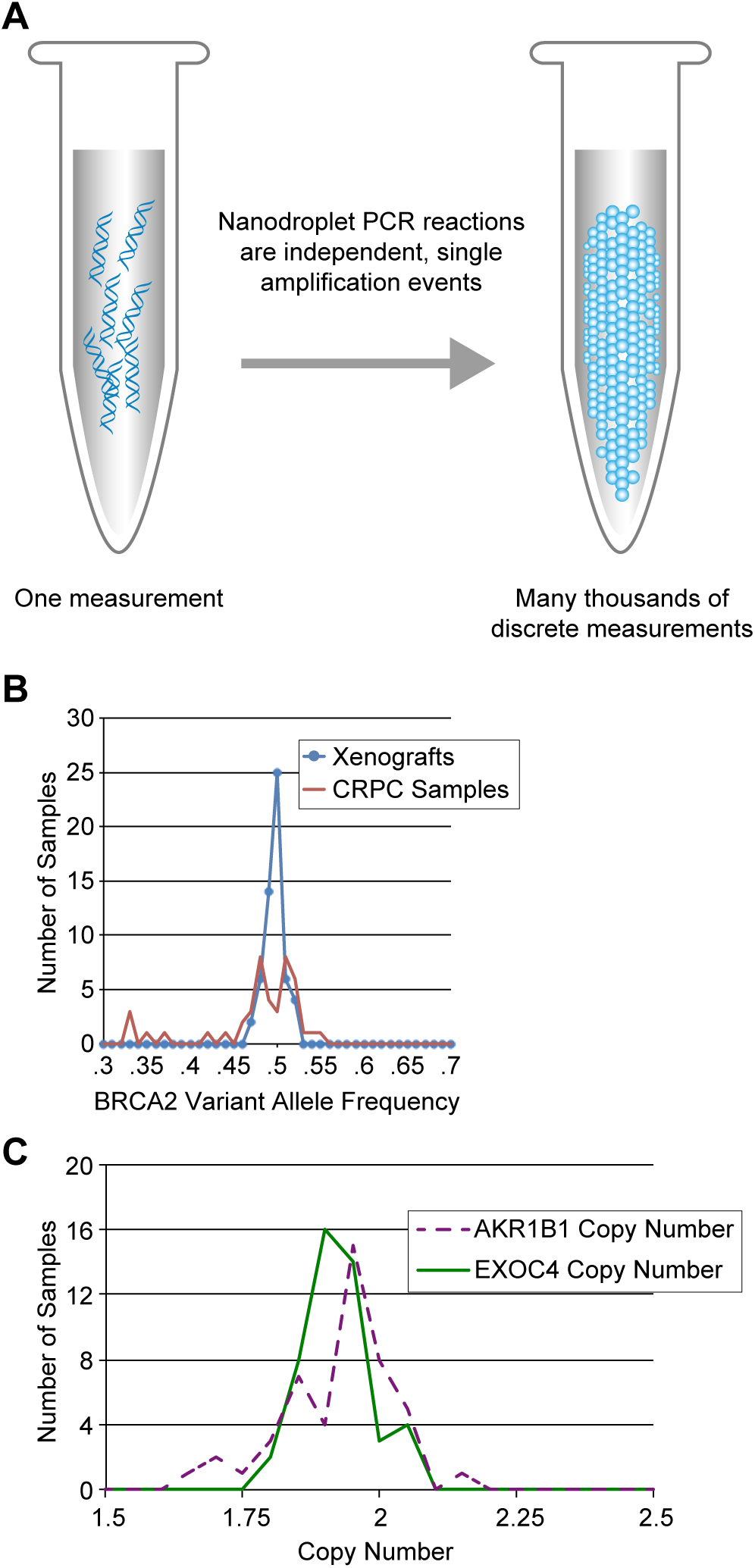
Assessment of mutations and copy number by droplet digital PCR. (A) ddPCR schematic. (B) Distribution of allele frequencies for the BRCA2 locus of Figure 1C as measured by ddPCR. Bulk tumor samples exhibit mean and standard deviation 0.500±0.011. Cultured cells exhibit mean and standard deviation 0.478±0.055. (C) Distribution of ddPCR-measured copy number at two loci expected to be diploid in TM00099.

As expected, ddPCR AF estimates were more precise and accurate compared to the panel sequencing. For example, for the heterozygous *BRCA2* mutation, ddPCR data across the bulk xenograft samples showed AF=0.50±0.011, improving on the exome-seq uncertainty (Figure 2B). We cultured conditionally reprogrammed progenitor cells (CRPCs) from the same PDX models, and these exhibited a wider distribution of AFs (0.478±0.055). The deviation of the CRPC samples from AF=0.5 was driven by outliers with AF=0.32-0.37, suggesting discrete changes consistent with population bottlenecks during *in vitro* culturing. CN measurements were robust and absolute. For example, two TM00099 control loci expected to be diploid from the exome-seq data exhibited ddPCR CN close to expectations across 47 samples (*EXOC4* CN=1.92±0.10; *AKR1B1* CN=1.92±0.06. See Fig. 2C). The *EXOC4* and *AKR1B1* copy numbers were not correlated (r=-0.19, p-val=0.21), indicating that the deviations were due to measurement uncertainties rather than sample-specific quality issues. CN measurements at separate loci within a gene were also highly correlated (two *LRP1B* loci: r=0.82, p-val=1e-12). Together the AF and CN data yielded subclone cellularity uncertainty <3% even from measurements at a single (diploid) locus.

### Symbiotic growth of recurrent subclones

ddPCR AF data provided evidence for both culturing-dependent bottlenecks and subclonal mixing in the TM00099 samples. For example, some loci (e.g. *BRCA2*) exhibited relatively stable AF values across samples while other loci exhibited a broader AF range (e.g. *ERBB4*) as shown in Fig. 3A. At stable loci, the variance in AF was dominated by culturing bottlenecks -- the bulk xenograft samples had *BRCA2* AF values centered at 0.5 (black); CRPC cultures grown from bulk were centered at 0.5 but with more outliers (blue); and CRPC colonies grown from single cells showed the most extreme outliers and at quantized values (red). In contrast, at *ERBB4* we observed a continuous range of AF values, with some of the lowest values coming from bulk xenograft samples. This continuous behavior suggested that the observed *ERBB4* values were the result of mixing of subclones of different genotypes, with any value for the mixing ratio allowed.

**Figure 3.**
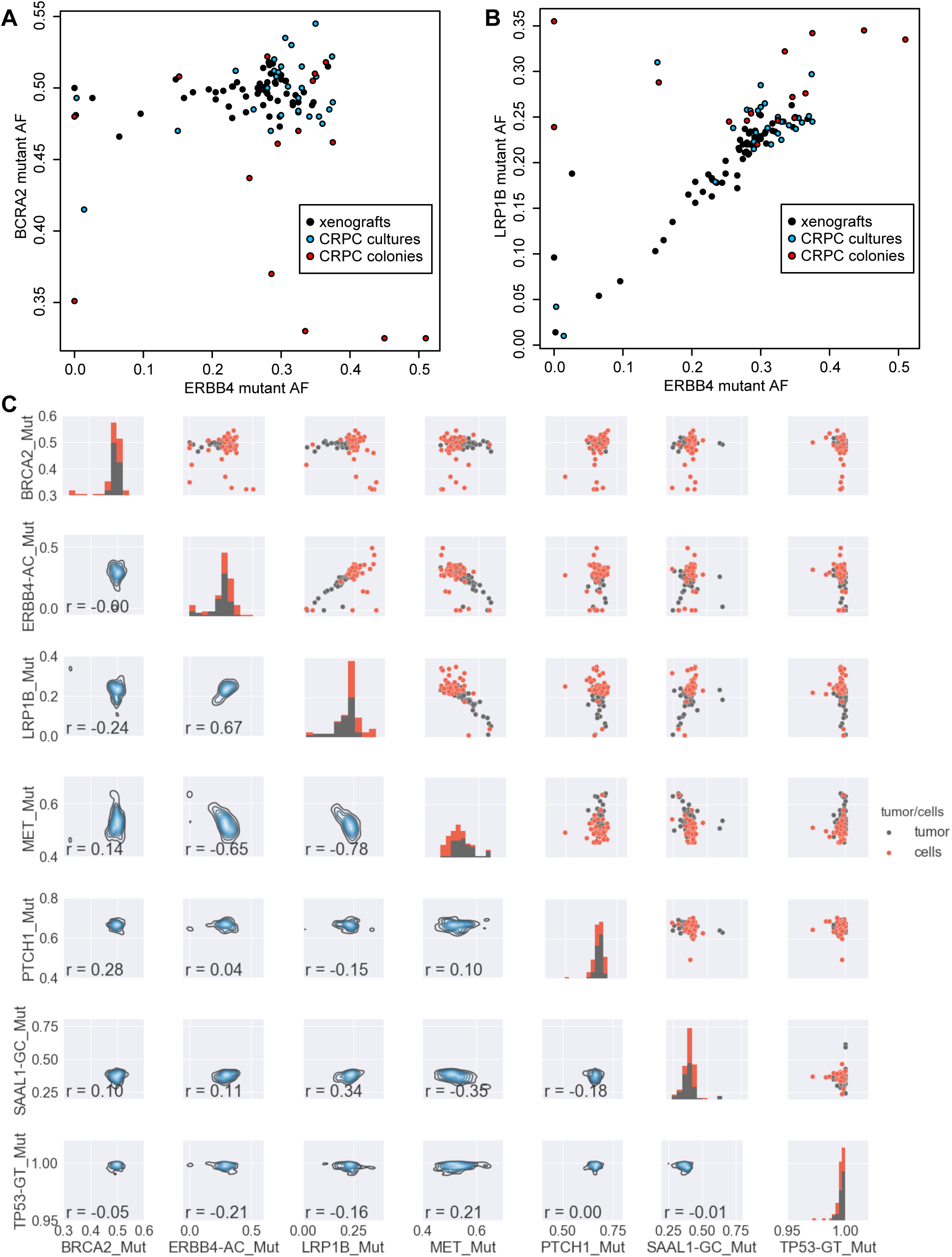
Comparisons of ddPCR mutant allele frequencies across samples. (A) Comparison of allelefrequencies of variants observed at the BRCA2 and ERBB4 loci in TM00099. Black dots are bulk tumor samples. Blue dots are conditionally reprogrammed progenitor cell (CRPC) cultures. Red dots are CRPC cultures established by growing from single cells. (B) Comparison of ERBB4 and LRP1B mutations. (C) (Upper right triangle): Pairwise plots of ddPCR-measured mutant allele frequencies for xenograft tumor samples (black) and CRPC samples (red). (Diagonal): Distribution of AF values for each mutation. (Lower left triangle): Density plots of AF values for pairs of mutations. Pearson correlations for each pair of mutations are also shown.

Comparison of other mutations also provided evidence for recurrent mixing between two subclones. For example, we observed a strong correlation between the AF values of *ERBB4* and *LRP1B* (r=0.67, p-value=2.6e-14) with dense occurrence of samples along the correlation line (Fig. 3B). These data can be explained by the presence of a subclone having both *ERBB4* and *LRP1B* mutations and another subclone lacking the two mutations, where each sample’s location along the Fig. 3B diagonal is determined by its ratio of the two subclones. Analysis of the full set of pairwise comparisons supported this conclusion (Fig. 3C). While several variants exhibited stable AFs consistent with truncal status in the tumor phylogeny (*BRCA2* ~0.5; *PTCH1* 0.6-0.7; *SAAL1* 0.3-0.4; *TP53* > 0.97), we observed an additional strong correlation between a mutation in *MET* and each of *ERBB4* and *LRP1B* (*MET* vs. *ERBB4* : r=-0.65, p-val=2.4e-13; *MET* vs. *LRP1B*: r=-0.78, pval=2.2e-16). The negative sign of the correlation indicated that the subclone that lacks the *ERBB4* and *LRP1B* mutations contains the *MET* mutation.

Copy number data further supported the conclusion of recurrent mixing between two subclones. We measured 9 loci in the TM00099 tumor, and the majority exhibited variance across samples larger than the ~0.1 CN measurement uncertainty observed for the control loci (Fig. 4A, control loci *AKR1B1* and *EXOC4* not shown). Similar to the AF data, we observed striking CN correlations between loci. In particular, pairwise comparisons within the group of *EPHA3*, *EPHA5*, *PTPRD*, *LRP1B,* and *ERBB4* and all had Pearson correlations r ≥0.73. We also observed strong anti-correlation between *AR* and the other loci, all with correlation values r ≤ −0.54. Like the *ERBB4* AF data, the CN data covered a smooth range of values, supporting their control by a continuous set of subclone mixing ratios. A heatmap of the correlations among all measured mutations and copy numbers is shown in Fig. 4B, indicating the strong associations between mutations and copy number events in the two subclones. Plots of all pairwise comparisons are given in Figure S5, and raw data are available in Table S1. A notable aspect of subclone 1 is its extremely high level of internal homogeneity. Within subclone 1 all pairwise correlations are r>0.69. In contrast, within subclone 2 *MET* AF and *AR* CNV are strongly correlated (r=0.71) but the other correlations are all ≤0.45. This indicates subclone 1 is more homogeneous than subclone 2, likely due to a recent population bottleneck.

The large number of samples allowed us to identify the genotypes of the subclones despite ubiquitous population mixing in each sample. We determined genotypes by extrapolating from a linear fit of AF values and absolute copy numbers, as illustrated in Figure 4C for *LRP1B* AF vs. CN (r=0.85, p-val=5.6e-14). These data lie close to a line connecting the points (CN=2, AF=0) and (CN=4, AF=0.25), indicating that subclone 1 has 1 mutant copy of *LRP1B* out of 4 total copies while subclone 2 has 2 wildtype copies of the *LRP1B* locus. We confirmed these inferences by measuring CN and AF in one sample expected to contain mostly subclone 1 (*LRP1B* AF=0.25, sample 1 in Table S1). For *LRP1B* we observed copy number number 3.6, approximately consistent with all cells in the sample having 1 mutant copy out of 4 alleles. Other loci also appeared to be homogeneous in this sample as their CN and AF values were plausibly consistent with all cells sharing a genotype, e.g. for *ERBB4* (AF=0.33 and CN=2.9: consistent with 1 of 3 alleles being mutant), *MET* (AF=0.48, CN=2.00: 1 of 2 copies are mutant), *PTCH1* (AF=0.66, CN=2.86: 2 of 3 copies are mutant), *TP53* (AF=0.99, CN=3.01: 3 of 3 copies are mutant), and *BRCA2* (AF=0.50, CN=4.01: 2 of 4 copies are mutant). This consistency further supported the conclusion of a recent population bottleneck for subclone 1.

**Figure 4.**
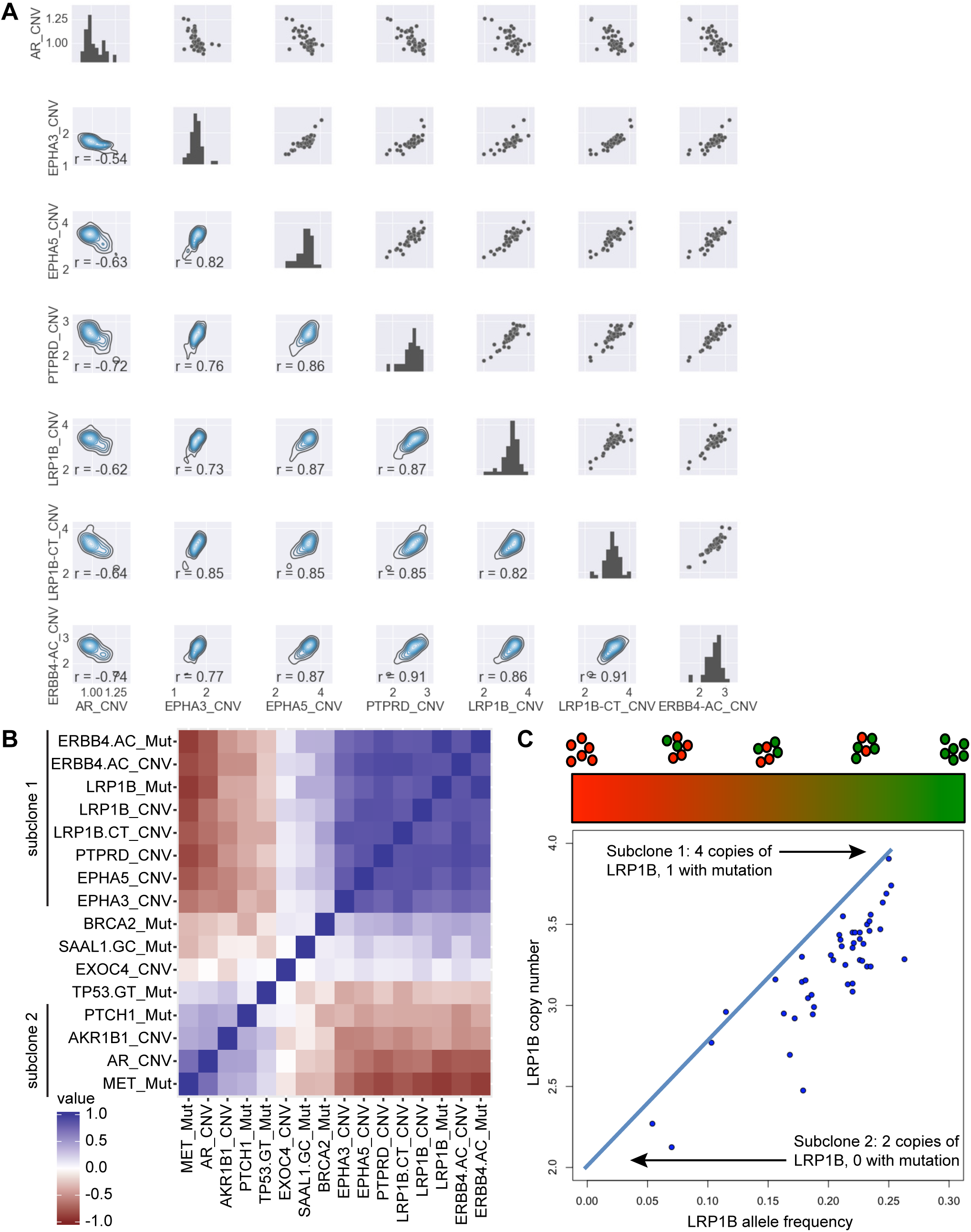
Resolution of subclones from ddPCR copy number and mutation measurements. (A) Pairwise plots of copy number variations in TM00099. Upper right triangle, diagonal, and lower left triangle are analogous to Fig.3C. (B) Pearson correlations among all ddPCR-measured mutant AFs and CNVs. The data comprise 47 TM00099 xenograft samples with data for all variables. (C) Comparison of LRP1B allele frequency and LRP1B copy number.

### Recurrent subclones are differentially selected by treatment

We next investigated whether the two subclones might differ in resistance to treatment. For this we analyzed ddPCR data from PDX residuals from the original exome-seq cohorts and a second set of xenograft cohorts treated for three weeks with cisplatin, cyclophosphamide, docetaxel, doxorubicin, or vehicle. Additionally, 6 mice that showed good response in the second cisplatin cohort were allowed to regrow without drug until they reached the pre-treatment starting volume of 150 mm^3^ (10-30 days) and were then retreated with a subsequent cycle of cisplatin before sequencing. ddPCR measurements were obtained for multiple xenografts from each cohort, allowing us investigate the relationship of tumor size and subclonal prevalence on a mouse-to-mouse basis.

ddPCR measurements indicated that the efficacy of cisplatin treatment was related to the post-treatment subclone distribution (Figure 5A). For the samples treated with cisplatin, *LRP1B* copy number, which increases with subclone 1, exhibited a strong correlation with tumor volume change after treatment (r=0.77, p-val =0.0008). This correlation was visible though statistically marginal when restricted to the samples given only one cycle of treatment (Figure 5A light green dots, r=0.51, p-val=0.19). The correlation was stronger for the samples allowed to regrow and be retreated for a second cycle (Figure 5A dark green dots, r=0.92, p-val=0.0092). The sample with the largest volume (Figure 5A green asterisk) was one subjected to one cycle of cisplatin and then regrown without further treatment, and it had the second highest value of *LRP1B* copy number. These data indicate that subclone 1 is more susceptible to cisplatin than subclone 2 but that in the absence of treatment subclone 1 grows faster. Neither *LRP1B* copy number nor tumor volume was correlated with the time at which the tumor was excised (Figure 5B, *LRP1B*: r=-0.17, p-val=0.54; Figure 5C %volume change: r=-0.25, p-val=0.36), i.e. tumor volume and relative subclone prevalence were each more impacted by individual-specific variations than by the total time or number of treatment cycles.

**Figure 5.**
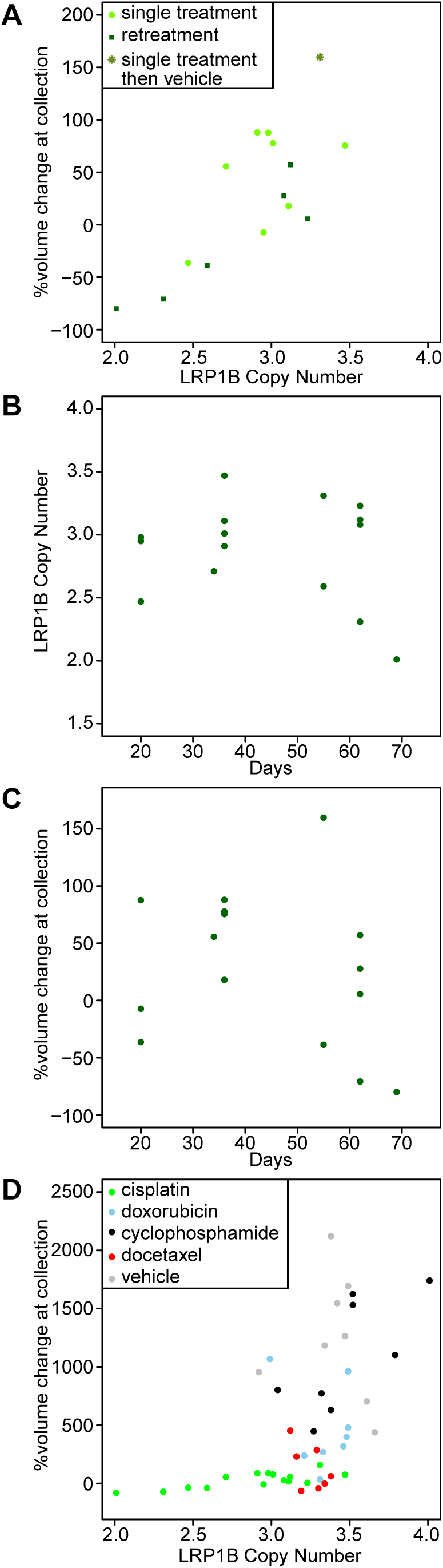
Relationship of intratumoral heterogeneity and tumor growth. (A) Comparison of LRP1Bcopy number and tumor volume after cisplatin retreatment. The y-axis indicates % volume change in the tumor at sample collection relative to the start of treatment. LRP1B copy number, a proxy for the relative cellularity of the two major subclones, is correlated with tumor volume change (r=0.77, p-val =0.0008). (B) Comparison of time vs. relative cellularity as quantified by LRP1B copy number. (C) Comparison of time vs. tumor volume during cisplatin treatment studies. (D) Comparison of tumor volume and LRP1B copy number for all treated TM00099 samples.

Comparison of treatments indicated that cisplatin was unique in selecting for subclone 2. The non-cisplatin cohorts exhibited larger *LRP1B* copy number values (ranging from 2.9-4.0) than the cisplatin-treated xenografts (2.0-3.5) as shown in Fig. 5D, providing evidence that cisplatin treatment selected for subclones with low *LRP1B* CN. For the non-cisplatin cohorts, correlations of subclone ratio vs. tumor volume were weaker. Cyclophosphamide-treated xenografts had a marginal correlation between volume change and *LRP1B* CN (r=0.71, p-val=0.05), and none of the other treatment cohorts exhibited significant correlations (doxorubicin r=-0.26, p-val=0.53; docetaxel: r=-0.54, p-val=0.20; vehicle: r=-0.17, p-val=0.68). Thus the two subclones have differential growth specific to cisplatin therapy, with a possible minor differential susceptibility to cyclophosphamide.

### Evolutionary bottlenecks occur commonly during growth

Although the dominant effect in the TM00099 data was the recurrent mixing of subclones 1 and 2, we also observed a minority of samples with outlier values not explainable by such mixing. Such outliers can only arise if population bottlenecks macroscopically increase the cellularity of a subclone with an alternate genotype. We identified outlier samples by comparing AF values to either a constant (*BRCA2, PTCH1, TP53, CSMD3*) or to the range of values possible from mixing of the subclones (*ERBB4, MET, SAAL1 -- LRP1B* AF was used as the predictive variable) (Fig. 6A; see Methods). 18 of the 101 samples exhibited outlier behavior along at least one the 7 AF dimensions, indicating that bottlenecks affect a significant minority of samples. Most samples had outlier behavior in only one dimension, though some samples deviated in up to 3 dimensions. As expected from culturing bottlenecks, a higher fraction of CRPC samples (25%) exhibited outlier behavior. However a non-negligible fraction of xenograft samples (12%) were also outliers. Different outlier samples were biased toward different cell populations, as we observed 12 distinct AF deviation patterns among the outliers (Fig. 6B). Thus these bottlenecks were not explainable by shared selection pressures for a small set of subclones across different samples.

**Figure 6.**
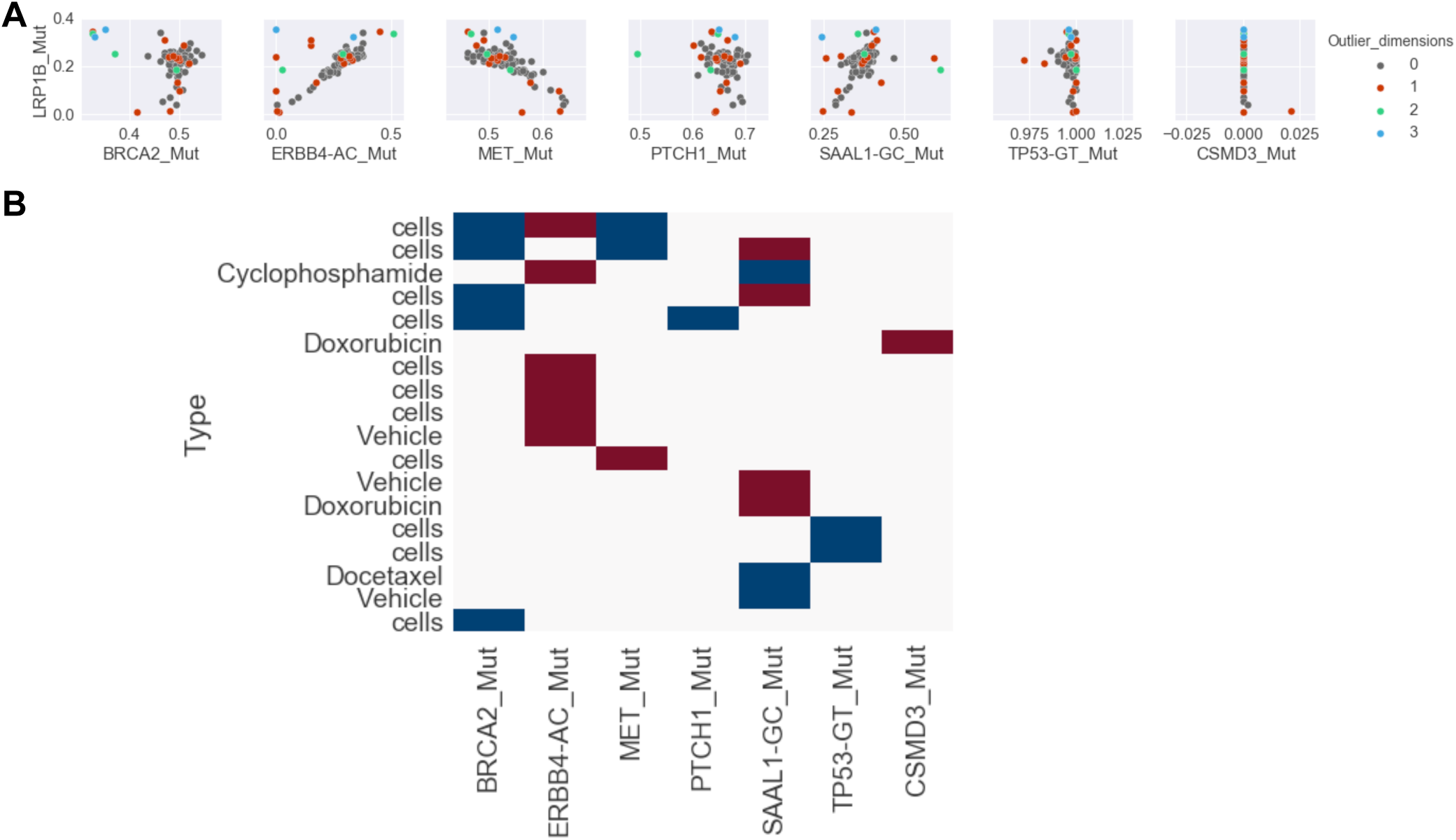
Samples with outlier allele frequencies. (A) Pairwise plots AF of 7 mutations vs. LRP1B AF.Samples with outlier behavior are denoted by red, green, and blue, where colors indicate the number of dimensions (i.e. AF values) in which each sample shows outlier behavior. (B) Samples with outlier behavior and the dimensions in which they deviate from expectations. Red indicates an excess AF value and blue indicates a deficient AF value. CRPC samples are labeled as “cells” and other labels indicate xenograft treatments.

To better understand the therapy-induced selection in the TM00099 model, we sought to compare how subclone distributions evolve under reduced selection pressures. However, the stable symbiosis of the two subclones suggested that there might be additional ecological constraints that impact TM00099 subclone evolution. As a simpler comparison we therefore interrogated the TM00096 tumor, for which our cancer panel data did not yield markers of commonly recurring subclones during treatment (Figure S6 and Table S1). For this analysis, we compared 8 untreated samples from the spatial dissection and propagation experiments (Figure 1D and Figure S3).

Analysis of ddPCR data at loci in the TM00096 samples revealed that spatiotemporally proximate sections have more similar AF values, consistent with expectations from spatially controlled genetic drift (Fig. 7A). For example, we measured AF values for mutations in *TTN, ERBB4,* and *DNMT3B*, which were detectable at low levels in many samples indicating they arose early in tumorigenesis (Figure S3). These mutations each spanned a range of AF values (*TTN*: 0-16.4%; *ERBB4*: 0-19.4%; *DNMT3B*: 0-2.3%) broader than the ddPCR measurement uncertainty (1.1%), and values were more similar for proximate samples. However, we also observed substantial differences in cellularity across samples inconsistent with Wright-Fisher drift (see Discussion). For example, a derived P3 sample contained a subclone with a *TTN* mutation in ~30% of its cells (assuming heterozygous diploidy; see Table S1), even though this subclone was undetectable in 4 of the other samples. Another P3 sample contained a subclone with an *ERBB4* mutation making up ~40% of its cells, even though AF was <3% in all other samples. Thus even in the absence of therapy we observed bottlenecks sufficient to replace more than one-third of the cells in a 20 mg fragment during a 2 month period.

**Figure 7.**
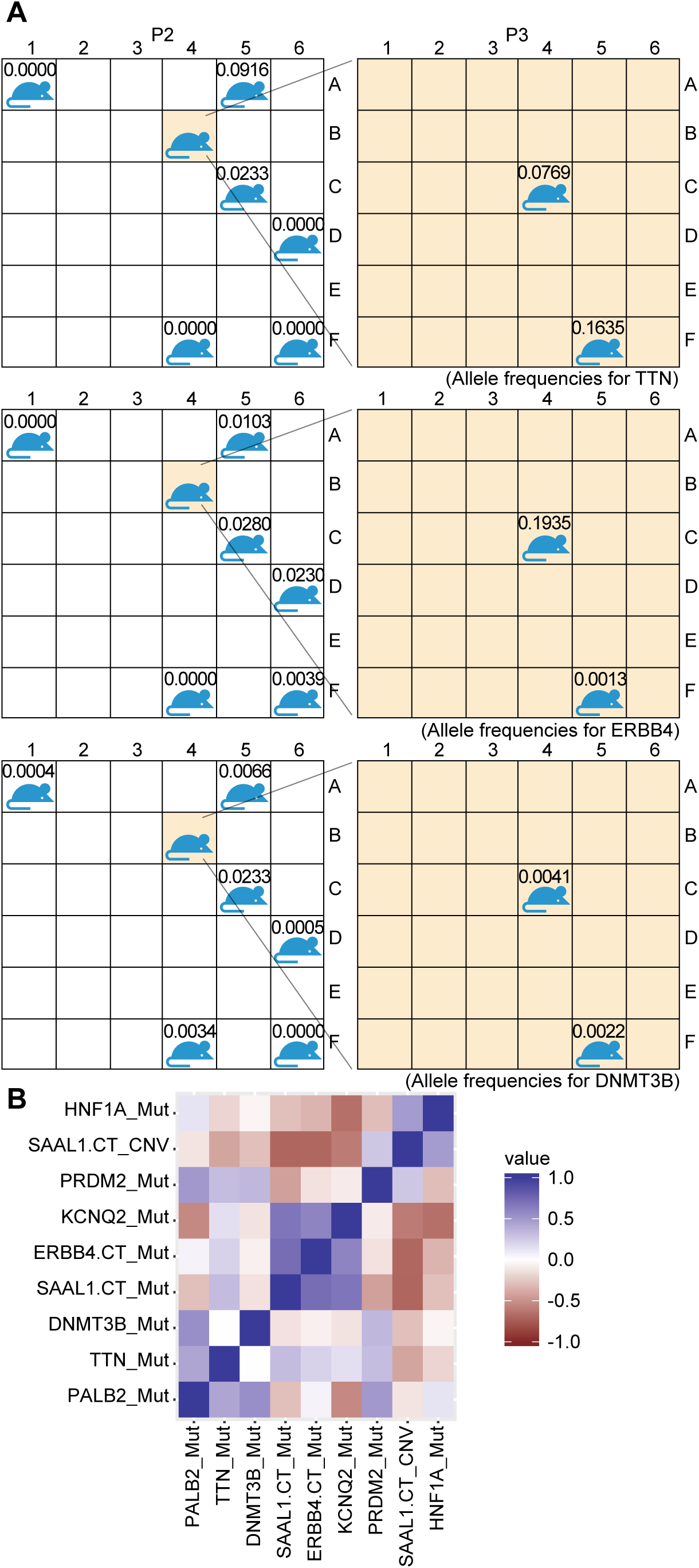
Spatiotemporal dependence of intratumoral heterogeneity. (A) Allele frequencies formutations in TM00096 P3 tumors grown from the P2 positions A1, A5, C5, D6, F4, and F6 (left) and a P3 grown from the P2 B4 positions dissected into 2 further samples P3 C4 and F5 (right). (B) Heatmap of correlations for ddPCR measurements of TM00096 spatiotemporal samples.

The evolutionary patterns among spatial samples were qualitatively different from those in the TM00099 treated samples. For the spatially dissected samples, no significant correlations were observed (p<0.05) between any pair of measured AFs or CNs (Figure 7B; see also Figure S6). In contrast, within each of subclones 1 and 2 in TM00099, every pair of markers was correlated with p<0.05. The lack of correlations within the spatial samples was consistent with the presence of multiple subclones whose cellularity levels varied independently.

## DISCUSSION

Using extensive xenograft and cell culture studies with high resolution ddPCR, we have presented to our knowledge the most exhaustive treatment-guided tumor subclonality deconstructions that have been performed in any tumor type to date. Importantly, ddPCR improved subclone quantification relative to the uncertainties that have been frequently described for exome sequencing approaches [28], allowing us to identify complex population evolution including symbiosis between recurrent subclones, frequent population bottlenecks, and differential drug sensitivity. Although multi-sample exome-seq combined with statistical inference approaches can improve subclone inference, such approaches are still limited by data quality, and we observed that ddPCR data improved PyClone subclone inferences over exome-seq (Fig S7). These studies demonstrate the value of precise copy number measurements when analyzing intratumoral heterogeneity from bulk measurements, providing a caveat to prior analyses that have relied on mutant allele frequencies alone [29].

### Potential drivers of subclone drug sensitivity

A major unexpected finding from the TM00099 tumor was that there are two stable populations that coexist, and we were curious what genomic differences might explain their differential cisplatin sensitivity. We analyzed if any mutant AFs or CN aberrations differed between treatment cohorts, and we found that all cohort pairs with a significant AF difference (Welch’s t-test, α=0.05, Figure S8) involved the cisplatin cohort, though none were significant after multiple testing (Benjamani-Hochberg) correction. However for CN comparisons, there were 9 significant differences after multiple testing correction, and 8 of 9 involved the cisplatin cohort. The two loci with the strongest differences between vehicle and cisplatin were *LRP1B* CN (average *LRP1B* CN across 2 measurements, p=0.00068) and *ERBB4* CN (p=0.00038). Direct comparisons with tumor volume also suggested that CN drives the differential sensitivity of the subclones (Table S2). Tumor volume was strongly correlated with the CN amplifications of subclone 1 -- *ERBB4*, *LRP1B*, *PTPRD*, and *EPHA5* CN all exhibited r ∈ [0.76,0.79] and p-value ∈ [4.8e-4, 9.4e-4]. In contrast, the strongest correlation between AF and volume was for *PTCH1* AF (r=-0.79, p-value=4.4e-4), though *PTCH1* AF varied little across samples (AF range=0.65-0.7) making it unlikely to be functionally important. Although our measurements cover only a limited set of the genome, these results suggest that copy number aberrations are more likely to distinguish the drug sensitivity of the subclones than point mutations, which may be related to the tumor’s positive tandem duplicator phenotype status [30].

### Tracing intratumoral population history

The large number of samples generated from diverse conditions allowed us to trace the population history within these tumors at unprecedented resolution, analogous to reconstructions of human population history. As illustrated by the correlations in Figure 4B, subclone 1 is internally more homogeneous than subclone 2, providing evidence that it is a more recently derived subclone. Thus the history of this tumor is similar to the human “Out-of-Africa” event [31] -- one subclone (subclone 1) emerged from a population bottleneck in the original population (subclone 2) and then expanded to make up a majority of the population. Interestingly, the absolute quantification of copy number in ddPCR provided additional evidence for this conclusion. As shown in Figure 4C, *LRP1B* has approximately 4 copies per cell in subclone 1 and approximately 2 copies per cell in subclone 2, showing that subclone 1 and subclone 2 are each relatively homogeneous at this locus. However, other loci behave differently. For *ERBB4*, a linear fit indicates that there are 2.8 copies on average in subclone 1 and 1.3 copies in subclone 2 (see Methods). Thus the subclone 1 value is slightly closer to an integer than the subclone 2 value. Because CN values must be integers in each individual cell, this suggests that subclone 2 is more heterogeneous than subclone 1, consistent with a population bottleneck that led to subclone 1. This effect, while modest, indicates how absolute CN data can reveal heterogeneity in a manner not accessible by relative CN measurements.

We also observed population bottlenecks that caused outlier subclonal distributions in 18% of all samples, including bottlenecks that distinguished spatially separated samples within the same xenograft. Surprisingly, even without therapy, small pre-existing cell populations were able to take over one-third of the cells of a ~20 mg region of a tumor in a period of only 2 months. It remains to be seen how much of these effects are due to selection versus drift processes such as spatial dispersal and neutral cell turnover. While the irregularity of the outlier populations supports the importance of drift [32], these allele frequency changes are too large to be explainable by neutral Wright-Fisher drift at the population sizes within xenograft fragments (~10^6^ in a 1mm^3^ fragment used for sequencing) and given the length of time (a few dozen cell replications). We speculate that these bottlenecks may be due to additional selection of subclones based on their epigenetics, and that the outlier populations may have more coherent epigenetic characteristics.

### Implications for dynamic treatment approaches

Finally, we consider the clinical importance of determining the population histories within these tumors (Fig. 8). In the TM00099 tumor, subclone 1 dominated the non-cisplatin xenografts (70% of cells on average, as interpolated from the *LRP1B* CN values) while the cisplatin-treated xenografts had reduced levels of subclone 1 (43% of cells on average). Thus therapy was more effective on the more recently-derived, faster-growing subclone. However, even during tumor shrinkage the more resistant population was difficult to kill completely. This corresponds to an ecology in which time-dependent treatment strategies such as metronomic therapy [33]or adaptive therapy [34] to extend the time of non-lethal tumor burden are suitable. For example, in adaptive therapy, treatment dosing is chosen based on dynamic changes in tumor volume, and this approach preferentially kills fast-growing sensitive populations while limiting takeover of the tumor by slow-growing resistant populations. Although ecologies susceptible to adaptive therapy ecologies have been synthetically constructed [35,36], to our knowledge ours is the first study to determine the genetics of such an ecology arising spontaneously *in vivo*.

**Figure 8.**
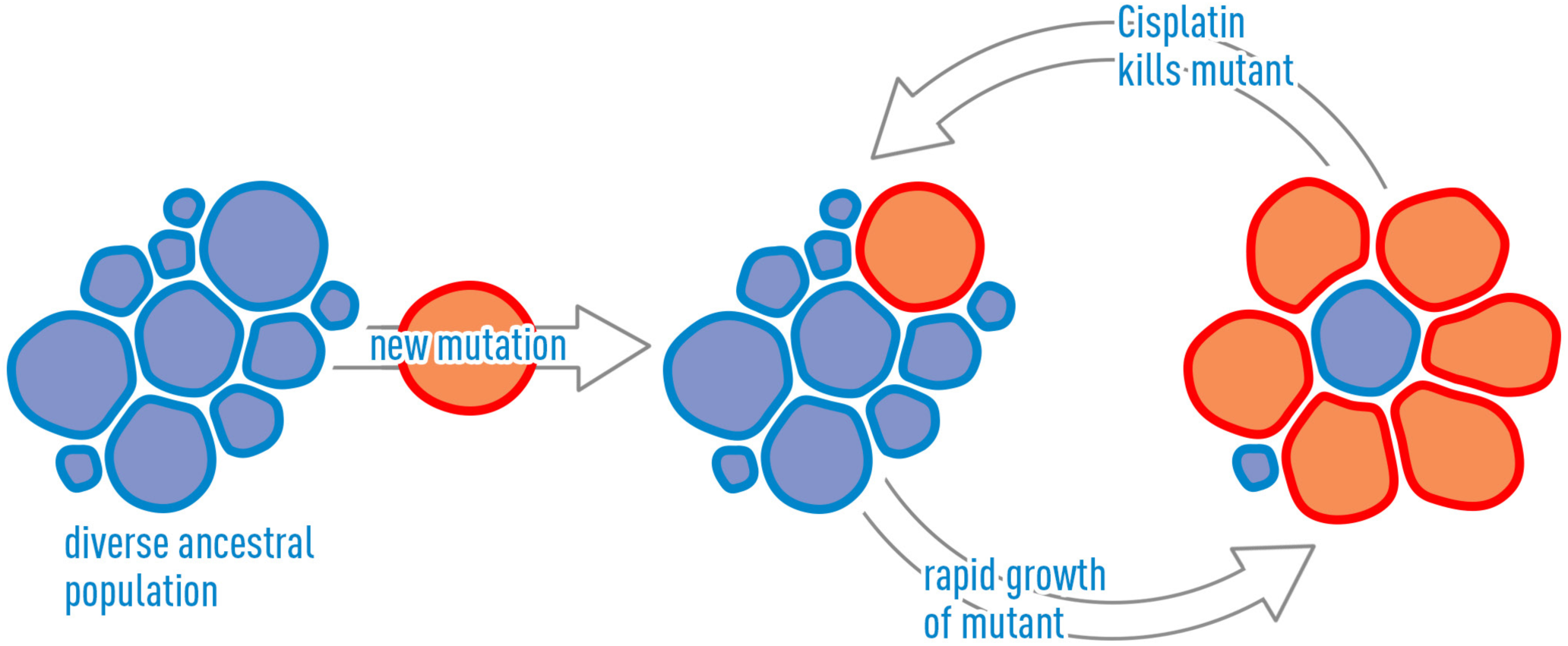
Schematic of evolutionary dynamics in the TM00099 tumor. Initially, the tumor contains adiverse ancestral population (blue), where diversity is indicated visually by the variability in cell sizes. A new mutation arises leading to a new population of cells (red). In the absence of treatment, this new population grows more rapidly, but it is also more susceptible to cisplatin treatment. Because the new population has arisen more recently, it is internally more homogeneous, indicated by the consistent sizes of the red cells. The relative growth behavior of the two populations makes the tumor susceptible to adaptive therapy cycles. The red and blue populations correspond to subclones 1 and 2, respectively, in Figure 4.

Recent reports have suggested that metronomic topotecan / pazopanib therapy is applicable to triple negative breast cancer [37] and other studies have recently demonstrated prostate cancer patients treatable by adaptive abiraterone therapy [38], making determination of the prevalence and genomics of such ecologies across cancer types a vital area for further inquiry. While there are non-negligible uncertainties in subclonal estimation for exome-seq-focused datasets such as TCGA [39], careful whole genome-based studies have shown that breast cancers often contain subclones with a high cellularity level [40]. Thus it is possible that many breast and other cancers have ecologies similar to ours, i.e. they contain high cellularity subclones whose relative proportions can be targeted for treatment. We note that, although we have focused our analysis on the cisplatin cohort from one patient, there may be mutations outside the exome panel that could distinguish balancing between common subclones in our other cohorts as well. Looking forward, we have seen that multi-sample data are essential for accurately distinguishing subclones, and circulating tumor DNA measurements may provide an approach to obtain such information for patients without the extensive xenografting efforts we have described here. ctDNA content has been suggested as a way to quantify tumor burden, but we speculate that quantitative subclonal decomposition from ctDNA may eventually have greater treatment relevance, as the observed ratio of subclones could be used in adaptive therapy mathematical models that stipulate treatment dose from tumor size [34].

An interesting last point is that the two subclones from the TM00099 tumor are likely to have symbiotic interactions with one another. We conclude this because all samples contained mixtures of the same major subclones, which is only possible if the two subclones are granularly intermixed. Moreover, the correlation between tumor volume and mixing ratio indicates that each xenograft must be relatively spatially homogeneous with respect to these subclones, otherwise randomness in the spatial locations of sequenced samples would have distorted the correlation. These observations suggest the presence of symbiotic interactions that synchronize the local mixing ratio of the two subclones. Previously, Marusyk et al [14] demonstrated ecological commensalism in a xenograft system generated from cell lines, making it plausible that there are commensal or mutualistic interactions in our system. The ratio of subclones does vary somewhat among the non-cisplatin samples (Figure 5C), but their composition is systematically different from the cisplatin samples, and both subclones continue to be present at non-negligible levels in the vast majority of samples. In the future, understanding of the dynamics and symbioses among naturally occurring subclones will benefit from finer mathematical modeling of selection and neutral evolution [41,42]. Such studies may reveal other ecological relationships that can be exploited to improve treatment strategies for cancer.

## MATERIALS AND METHODS

### Tumor Samples

For model TM00096, the tumor sample was obtained from a 45-year-old female patient with triple negative breast cancer subtype determined by immunohistochemistry. For this patient, anthracyclines and taxanes had been used as chemotherapeutic agents prior to surgery. The sample was obtained from a metastasis of the breast cancer to the patient’s lung. To establish the PDX, a fragment of the patient’s tumor was engrafted into a Nod scid gamma (NSG) mouse until stable growth, forming the P0 passage. A subsequent set of P1 mice xenografts were formed by dissecting the derived P0 tumor and reimplanting these into new mice. Fragments of a grown P1 passage were then used to begin the experiment in the P2 passage. During dissection of the P2, all fragments were weighed. For model TM00099, a similar procedure was used to establish xenografts. The tumor sample was obtained from a 44-year-old female patient with triple negative breast cancer diagnosed as invasive ductal carcinoma. The patient was treated with neoadjuvant taxol prior to resection, and the sample used for xenografting was obtained from the primary malignancy.

### Xenograft spatial dissection and temporal propagation

The patient-derived xenograft triple negative breast cancer tumor (JAX PDX resource model TM00096) from the P1 passage was dissected and a fragment (5-10 mm^3^) was implanted subcutaneously by trocar in the right flank of a female NOD.Cg-*Prkdc^scid^ Il2rg^tm1Wjl^*/SzJ (NOD scid gamma or, NSG) mouse. This P2 tumor was then allowed to grow for 118 days at which point its volume was >2000 mm^3^, after which it was spatially dissected into a 6x6 grid. Grid locations were annotated alphanumerically (A1 … F6). 6 of the P2 grid fragments (A2, A6, B5, D3, F1, F5) were sequenced via cancer panel sequencing. The total weight of the 36 P2 fragments was 883 mg. Fragments specified in Figure 1D were implanted into P3 mice. Implanted fragment weights ranged from 10.3-45.6 mg. After 58 days, four of these P3 xenografts (each with tumor volume > 1,000 mm^3^) were sacrificed. These were the xenografts generated from original positions B4, D6, F4, and F6 in the P2 tumor. Xenografts generated from the D6, F4, and F6 fragments were sequenced. We further dissected into a grid the P3 tumor that had been generated from the B4 fragment of the P2. Two spatial fragments corresponding to positions C4 and P5 in the P3 tumor were analyzed and sequenced. In addition, 3 other P3 xenografts were harvested at 72 days after engraftment. These were the P3 xenografts generated from fragments at positions A1, A5, and C5 in the P2 tumor.

### Xenograft treatment experiments

TNBC PDX models were established at The Jackson Laboratory In Vivo Services campus and tested for cisplatin, docetaxel, doxorubicin, and cyclophosphamide sensitivity. Briefly, patient tumor material acquired from biopsy or surgical resection was implanted subcutaneously into the flank of NOD-scid IL-2r gamma-chain null female mice (8–10 weeks old). Models were considered established when log-phase growth in a second passage was evident. Individual tumor-bearing mice were randomized into treatment cohorts of at least six animals each on an accrual basis when tumors reached a volume of 150 mm^3^ (day 0), at which point each tumor model was assessed for its response to treatment. The treatments used for each drug were: Doxorubicin – once weekly, 2 mg/kg iv, for 3 weeks; Cyclophosphamide – once weekly, 40 mg/kg iv, for 3 weeks; Docetaxel – once weekly, 10 mg/kg iv, for 3 weeks; Cisplatin – once weekly, 2 mg/kg iv, for 3 weeks. Changes in tumor volumes were measured using digital calipers for 4 weeks from the beginning of the treatment or until tumor volumes reached the 1,500 mm^3^ end point. For the retreatment study, TM00099 and TM00096 mice cohorts were established and subjected to a first cycle of treatment as described. However, instead of being harvested at the end of the treatment, tumors were maintained in their host mice and allowed a drug-holiday period until they grew back to their original pre-treatment volume (~150 mm^3^), at which point a second cycle of treatment was administered following the same protocol described above.

### Conditionally Reprogrammed Progenitor Cells

We cultured Conditionally Reprogrammed Progenitor Cells (CRPCs) in vitro from PDX tumors adapting an approach published by Yuan et al [43]. CRPC cultures are heterogeneous, and single cell clonal lines can be derived from them. Tumors from PDX models were dissociated into single cells with collagenase and cultured on irradiated 3T3J2 mouse feeder cells in media to form isolated epithelial colonies. Individual colonies were harvested and expanded as single cell clonal lines (red dots in Figures 3A and 3B). We also cultured some PDX tumor samples analogously on feeder cells but without separating into single cells, yielding bulk samples propagated in the in vitro environment. These correspond to the culture samples (blue dots in Figures 3A and 3B).

### Targeted exome sequencing panel

The Jackson Laboratory cancer panel is a targeted exome sequencing panel consisting of 358 genes. Inclusion of a gene on the panel was based on a three-tier criteria ladder, which consists of genes targeted by a drug (tier 1), genes having only causality in cancer but not drug targeted (tier 2), and genes only implicated in cancer as determined by Jackson Laboratory researchers (tier 3). Ten-percent (41/358) of the genes on the panel are identified as causal, 53% (172/358) are targeted by drugs and/or causal, and the remaining 37% (145/358) are research grade. Genes in tiers one and two were further annotated to signaling pathways involved in efficacy of targeted therapies using the Rat Genome Database Pathway Ontology. The highest percentages of genes on the panel reside in *EGFR*, *MAPK*, *mTOR*, and *p53* signaling pathways. The full list of genes and loci is available in Table S3.

### Mutation calls from cancer panel

FASTQ files were analyzed by a custom pipeline that includes tools to assess read quality, alignment, and variant calling. Human and mouse reads were separated using Xenome [44], after which only human reads were considered for further analysis. Human reads were quality trimmed and filtered using the NGS

QC toolkit to remove reads with > 30% low quality bases (Q < 30) [45]. The resulting reads were aligned to the human reference genome (hg19) using BWA [46]. Duplicates were removed using Picard (http://picard.sourceforge.net) and the resulting alignments were processed to minimize artifacts by realigning around indels and recalibrating the base quality score using the Genome Analysis Tool Kit (GATK) software suite [47]. Single nucleotide variants (SNVs) were called using GATK with HaplotypeCaller in GVCF mode [47], and stored to variant call format (VCF) [48] files. Variants of low quality or allele frequency <0.01 were removed, while variants with low sequencing depth (<140) were flagged. Genes belonging to the Mucin gene family (*MUC16*, *MUC17*, *MUC4*, *MUC5B*) were removed from the heterogeneity analysis, since they contain variable number tandem repeats making them polymorphic even in normal genomes. Common dbSNP variants were removed to filter germline effects. The dbSNP definition of a common variant is one polymorphic in at least one population in the 1000 Genomes project [49], having minor allele frequency >0.01, and present in at least two 1000 Genomes samples [50].

For the spatial and temporal analyses of cancer panel data, we performed mutation calling with MuTect [51] using the P0 (the first stable xenograft after implantation from the patient tumor fragment) as control. MuTect using the vehicle as control gave very similar results (data not shown). For these analyses, the MuTect pipeline was chosen over the GATK pipeline because it provided improved sensitivity for detecting low AF mutations.

### CNV calls from cancer panel

Exon level copy number profiles from the cancer panel were estimated using CONTRA [52]. The statistical significance of segment copy number levels were recalibrated using ConReg-R to improve the false discovery rate estimates [53]. We established a "normal" baseline comprised of 3 unrelated HapMap samples: NA12877, NA12878, and NA18507). These HapMap samples were sequenced as controls to minimize depth and sequence-related biases from cancer panel sequencing. The stability p-value of each exon-level copy number was defined based on the standard deviations of 20 adjusted log-ratio values obtained from 20 bins for each exon. When an exon was too small to divide it into 20 bins, a smaller number was used based on the constraints: minimum read depth=50, minimum bin width=20 nt. Deletion or amplification was determined based on the behavior of all bins for each exon. Additionally for the TM00096 sample we compared to copy number computed using ASCAT [54] from Affymetrix SNP6.0 array data of the tumor P0 passage. Similar to the CONTRA analysis, we observed that subclonal copy number inferences under this method were parameter-sensitive.

### Detection of copy number variation and gene mutations by ddPCR

The QX200 Droplet Digital PCR (ddPCR) System (Bio-Rad Laboratories) was used for detection of copy number variation (CNV) and/or gene mutations. Human gene-specific primers and probes were designed based on the human genome reference hg19 (GRCh37). In this study, *CUL1* was used as a reference gene for CNV assays since its absolute copy numbers were close to 2.0 for models TM00096 and TM00099. Absolute copy numbers were estimated by Affymetrix SNP6.0 array for P0 samples of TM00096 and TM00099. We selected mutations for which allele frequencies were variable by treatment but relatively stable for control mutations. For point mutations, we designed the primers and a FAM-labeled probe that targeted the mutation in the target gene as well as a HEX-labeled probe that targeted the wild type allele, respectively. ddPCR reactions were performed according to the manufacturer’s protocol. Briefly, 10ng DNA template was mixed with the PCR Mastermix, primers, and probes to a final volume of 20 μL, followed by mixing with 60 μL of droplet generation oil to generate the droplet by the Bio-Rad QX200 Droplet Generator. After droplets were generated, they were transferred into a 96-well PCR plate and then heat-sealed with a foil seal. PCR amplification was performed using a C1000 Touch thermal cycler and once completed, the 96-well PCR plate was loaded on the QX200 Droplet Reader. All ddPCR assays performed in this study included two normal human controls (NA12878 and NA10851) and two mouse controls (NSG and XFED/X3T3) as well as a no-template control (NTC, no DNA template). All samples and controls were run in duplicates. Data was analyzed utilizing the QuantaSoft™ analysis software.

Fractional abundance (mutation) or copy number of the duplicate wells was calculated and a t-test was performed to determine statistical significance compared to the controls.

### Estimation of copy number values in subclones

Based on the logic of Figure 4B, we estimated the copy number at *ERBB4* in subclones 1 and 2. To do this, we used all samples with measurement data and performed a linear regression of *ERBB4* CN versus *LRP1B* allele frequency. The estimated value for *ERBB4* CN in subclone 1 was the linear interpolation at *LRP1B* AF=0.25. The estimated value for CN in subclone 2 was the linear interpolation at *LRP1B* AF=0. See also Figure S5.

### Outlier detection

For the TM00099 mutations not impacted by mixing of the two major subclones (*BRCA2, CSMD3, PTCH1, TP53*), we identified outlier AF values by calculating a median-centered z-score for each sample: z(gene, sample) = (AF(gene, sample) – Median AF(gene, all samples)) / σ(gene, all samples). Samples with |z| > 3 were considered outliers. For the variables impacted by subclonal mixing of the two major subclones (*ERBB4, MET, SAAL1*), we calculated outliers based on their perpendicular Euclidean distance D to the line determined by each variable plotted versus *LRP1B* AF. These lines were those connecting the pairs of points: (*LRP1B, ERBB4*): (0,0) and (0.25,0.33); (*LRP1B, MET*): (0,0.67) and (0.25,0.5); (*LRP1B, SAAL1*): (0,0.25) and (0.25,0.4), respectively. A sample was considered to be an outlier for D> 0.07. This is much larger than the ddPCR AF measurement uncertainty of 0.011 (Figure 2B).

### Ethics Statement

The animal care rules used by The Jackson Laboratory are compatible with the regulations and standards of the U.S. Department of Agriculture and the National Institutes of Health. The protocols used in this study were approved by the Institutional Animal Care and Use Committee (IACUC) of The Jackson Laboratory under protocol number 12027. Mice used in this project were euthanized by carbon dioxide asphyxiation in a manner consistent with the 2013 recommendations of the American Veterinary Medical Association Guidelines on Euthanasia.

## ACKNOWLEDGMENTS

Research reported in this publication was supported by the National Cancer Institute under award numbers P30CA034196, R21CA191848, and U24 CA224067. The content is solely the responsibility of the authors and does not necessarily represent the official views of the NIH. JHC also acknowledges support from the Hope Foundation. The authors would like to thank Jason Li for valuable discussions about CONTRA. The authors thank Quaid Morris, Shankar Vembu, and Wei Jiao for discussions about tumor phylogenetic decomposition. The authors thank Jane Cha, Zoe Reifsnyder, and Matt Wimsatt for assistance with visualizations. We also thank scientific research services at The Jackson Laboratory including Doug Hinerfeld, Vanessa Spotlow, Janet Pereira, Bill Buaas, and Anuj Srivastava.

## SUPPORTING INFORMATION CAPTIONS

**Figure S1.** A high specificity mutation-calling pipeline for comparative evolutionary analysis across exome-seq samples.

**Figure S2.** Sanger sequencing results across multiple samples for the ERBB4 gene. Sample G1 does not have the ERBB4 mutation, while Samples G2-G5 show low levels of it. These samples were obtained from residuals of the treated TM00099 tumors, though not the same samples used for the exome-seq.

**Figure S3.** Spatiotemporal dissection of TM00096 tumor. (A) Fragments from the dissected P2 xenograft (location specified by rectilinear coordinates) were engrafted into P3 mice, grown, and sequenced. One P3 was dissected and two fragments (C4, F5) were sequenced (bottom row). (B) Visualization of ddPCR mutant AFs. Darker shading indicates higher AF, with mutations distinguished by wedge angle. (C) Alternate visualization with mutations in separate plots. (D) Inferred phylogenetic tree. HNF1A, KCNQ2, SAAL1-CT, and PALB2 have high AF in all samples, indicating truncal occurrence. DNMT3B, TTN, and ERBB4 are in multiple samples but often at low AF(<3%), indicating early but persistent subclonality.

**Figure S4.** Exome-seq copy number uncertainties for TM00096. CONTRA-based CN estimates are plotted vs mutant AF at the MUC4 locus. Correlation of CN and AF suggests importance of CN to cellularity estimation. However, CN estimates vary substantially and do not correspond to integer values, limiting inference of subclone genotypes and cellularity levels.

**Figure S5.** Pairwise comparisons of all mutant AFs and CNVs for the TM00099 tumor. Only samples that had sufficient DNA to samples to perform measurements at all loci are shown. Full data are in Table S1.

**Figure S6.** Pairwise comparisons of ddPCR measured quantities for TM00096. Green = spatiotemporal samples. Blue = treatment samples. The spatiotemporal and treated samples form two clusters because the spatiotemporal samples have a recent common ancestry.

**Figure S7.** Comparison of Pyclone results for panel exome sequencing and ddPCR. (left) The posterior similarity matrix generated by PyClone based on variant AF (allele frequency) and CN (copy number) estimates for all TM00099 samples as measured on the CTP exome panel. AFs of variant loci were computed by GATK, while copy numbers were estimated by CONTRA. Mutations called by the pipeline of Figure S1 are shown. (right) The posterior similarity matrix generated by PyClone with variant AF and CN estimates for TM00099 samples for sites measured by ddPCR. The improved inference from ddPCR is apparent from the stronger bimodality in correlation values. Pyclone was run with option –prior total_copy_number and read counts were modeled using the pyclone_beta_binomial option.

**Figure S8.** Comparisons of mutant AF and CN across single treatment cohorts. (A) Significant mutant AF comparisons (Welch’s test, alpha = 0.05, no multiple hypothesis correction). (B) Chord diagram of cohorts with significant mutant AF differences. Line width indicates p-value. Average cohort AF is indicated under each gene and by pink scale. (C) Significant CN comparisons (Welch’s test, alpha = 0.05, Benjamini-Hochberg correction). (D) Chord diagram showing significant CN differences.

**Table S1. ddPCR data.** Tables of all ddPCR and tumor volume data for samples derived from patient models TM00099 (sheet 1) and TM00096 (sheet 2).

**Table S2. Potential driver correlations.** Correlations between individual markers and tumor volume for all cisplatin-treated samples.

**Table S3. Cancer panel loci.** List of loci in exome cancer panel.

